# Illumina and Nanopore methods for whole genome sequencing of hepatitis B virus (HBV)

**DOI:** 10.1101/470633

**Authors:** Anna L McNaughton, Hannah E Roberts, David Bonsall, Mariateresa de Cesare, Jolynne Mokaya, Sheila F Lumley, Tanya Golubchik, Paolo Piazza, Jacqueline B Martin, Catherine de Lara, Anthony Brown, M Azim Ansari, Rory Bowden, Eleanor Barnes, Philippa C Matthews

## Abstract

Advancing interventions to tackle the huge global burden of hepatitis B virus (HBV) infection depends on improved insights into virus epidemiology, transmission, within-host diversity, drug resistance and pathogenesis, all of which can be facilitated by the large-scale generation of full-length virus genome data. Here we describe advances to a protocol to exploit the circular HBV genome structure, using isothermal rolling-circle amplification to enrich for HBV DNA and to generate concatemeric amplicons containing multiple successive copies of the same genome. We show that this product is suitable for Nanopore sequencing as single reads, as well as for generating short-read Illumina sequences. Nanopore reads can be used to implement a straightforward method for error correction that reduces the per-read error rate, by comparing multiple genome copies combined into a single concatemer and by comparing reads generated from plus and minus strands. Thus we can achieve improved consensus sequencing accuracy of 99.7% and resolve intra-sample sequence variants to form whole-genome haplotypes. The combination of isothermal amplification and Nanopore sequencing offers the longer-term potential to develop point-of-care tests for HBV, which could also be adapted for other viruses.

## Introduction

Chronic hepatitis B virus (HBV) infection affects an estimated 250-290 million individuals worldwide, resulting in around 800,000 deaths from chronic liver disease and hepatocellular carcinoma each year ^1,2^. The status of HBV infection as a globally important public health problem is highlighted by United Nations Sustainable Development Goals, which set a target for HBV elimination by the year 2030 ^3^. An improved understanding of the molecular biology, epidemiology, infection dynamics and pathophysiology of HBV is a crucial step towards reducing the global burden of HBV disease. Despite the availability of a robust prophylactic vaccine and safe suppressive antiviral therapy, HBV has remained endemic - and neglected - in many populations ^4^. Large-scale virus genome sequencing to provide more complete genetic information at the population and individual level can shed light on the limitations of current interventions ^5^, and inform new strategies for elimination. New, large-scale sequencing initiatives require improved methodologies that are efficient, accurate, sensitive and cost-effective.

In the context of clinical and public health settings, HBV sequencing can provide information that is useful in characterizing virus genotype, potential transmission networks, drug and vaccine resistance, and aspects of the dynamics of infection ^5–7^. Traditional Sanger sequencing can derive consensus sequences (usually of sub-genomic fragments), and next-generation technologies such as Illumina can interrogate within-sample diversity at the whole-genome level. Sequencing complete virus genomes at depth, while also preserving mutation-linkage information (ie. complete haplotypes), remains an important goal that will lead to more accurate phylogenetic characterisation of viral quasispecies within infected hosts, which can in turn be interpreted to study virus transmission and the evolutionary dynamics of drug and immune escape.

‘Third generation’ (i.e. single-molecule) sequencing approaches including those based on nanopores (Oxford Nanopore Technologies, ONT) ^8,9^, have the potential to revolutionise virus genome sequencing in several ways. First, they can produce genome-length reads that encompass all of the mutations within a single virus particle. Second, Nanopore technology is portable and provides sequence data in real time, potentially enabling sequencing as a point-of-care test. However, Nanopore sequencing has been adopted with caution because of its high raw error rates ^10^. While error-corrected Nanopore consensus sequence may be sufficiently accurate for many uses, raw-read accuracy remains a concern if it is to be used for the assessment of within-sample (between-molecule) diversity. One strategy to reduce error rates from single source molecules is to create concatemeric (chain-like) successive copies of each template, so that a single concatemer contains several reads of each base from the original molecule. This approach has been demonstrated in the circularization of 16S bacterial DNA sequences followed by ‘rolling circle amplification’ (RCA) using a high-fidelity DNA polymerase ^11^.

HBV has an unusual, circular, partially double-stranded (ds) DNA genome of 3.2kB ^12^. The combination of double and single stranded DNA could cause technical problems for sequencing, since library preparation methods are usually specific for either single- or double-stranded DNA templates. HBV isolates have previously been sequenced with Nanopore technology ^13^ using a full-length genomic PCR approach to enrich for HBV sequences. Whilst this approach worked well for the two samples tested, the high error rate observed in this previous paper (~12%) could only be corrected at the consensus level, meaning the data could only be used to determine the majority strain in the sample. In this study we build on a published method for HBV enrichment and amplification from plasma ^14–16^, which generates intermediates that are suitable for sequencing by Nanopore or Illumina. We implement novel analytical methods to exploit concatemeric reads in improving the accuracy of Nanopore sequencing of HBV for use in research and clinical applications.

## Results

### Completion ligation and rolling circle amplification prior to Illumina sequencing of full-length HBV genomes

We applied sequencing methods (as shown in Fig 1) to plasma from three different individuals with chronic HBV infection (Table 1). We first set out to convert the partially dsDNA viral genome to a complete dsDNA HBV molecule using a completion-ligation (CL) method ^15^, so that sequencing libraries could be generated using kits that require dsDNA as input. Following CL, genomes were amplified by rolling circle amplification (RCA) ^14,15^. We confirmed an increase in HBV DNA after RCA by comparing extracted DNA to RCA products using qPCR (for methods, see Suppl Methods 2). Using DNA products derived from CL alone and from CL followed by RCA, we prepared sequencing libraries and sequenced them using an Illumina MiSeq instrument.

**Fig 1:**
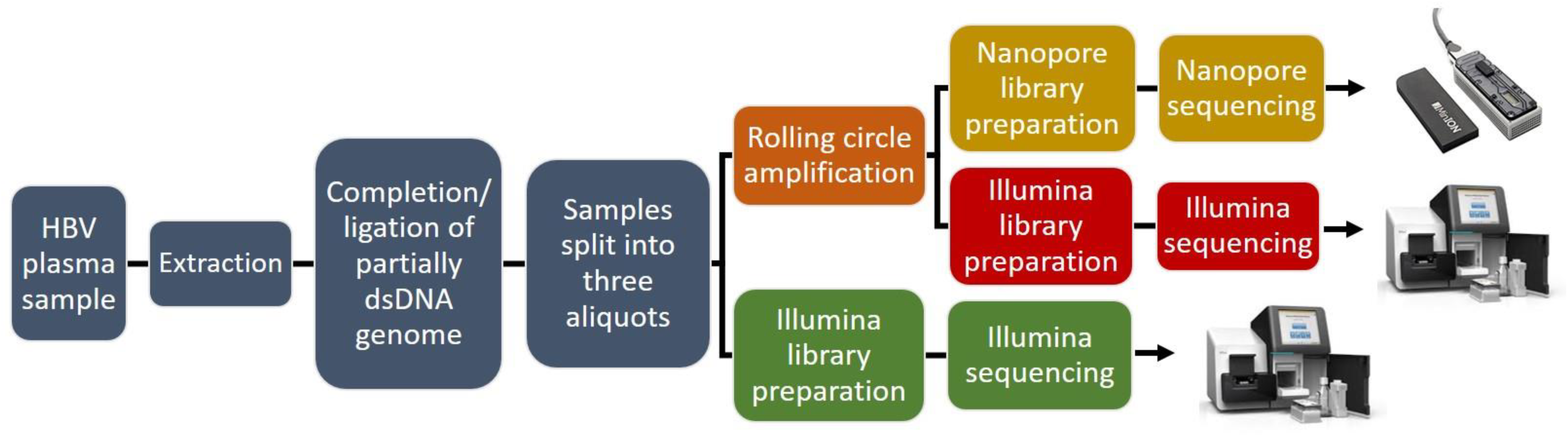
Flow diagram to show pathway of sample processing from plasma through to HBV genome sequencing on Nanopore (yellow) and Illumina (red and green) platforms.

**Table 1:**
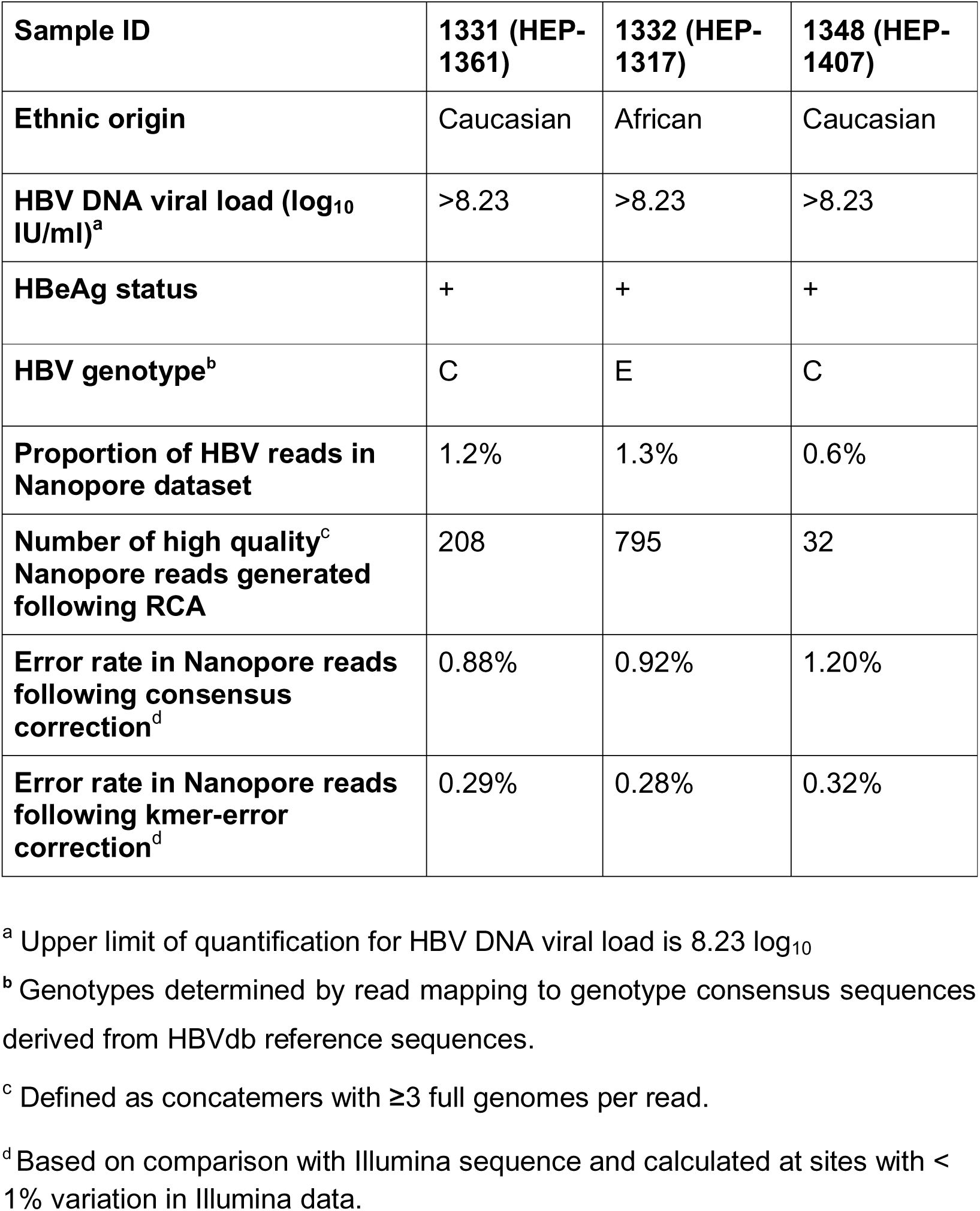
Details of samples used for HBV sequencing. Patients were adults with chronic HBV infection, enrolled through a cohort in Oxford, UK.

Both the CL and CL+RCA methods generated reads that covered the whole HBV genome for all three samples (Fig 2A). The relative drop in coverage across the single-stranded region of the HBV genome disappeared after RCA, suggesting a preferential amplification of intact whole genomes. We also observed a region of reduced coverage corresponding approximately to nt 2500-2700, in all samples. Further examination of the sample with the sharpest drop in coverage across this region (sample 1348) revealed a drop in the density of insert ends in the region (Suppl Fig 1) and resultant disruption to insert size (Fig 2B), consistent with inefficient digestion by the Nextera transposase. Reasons for the reduced coverage are unclear; no nicks in the HBV genome have been described in this region, but there may be some secondary structure present. GC content may also be a contributing factor: GC bases in the region nt 2500-2700 account for 35-37.5% in the Illumina consensus sequences, in contrast to the rest of the genome, where GC content is 48-49.5%.

**Fig 2:**
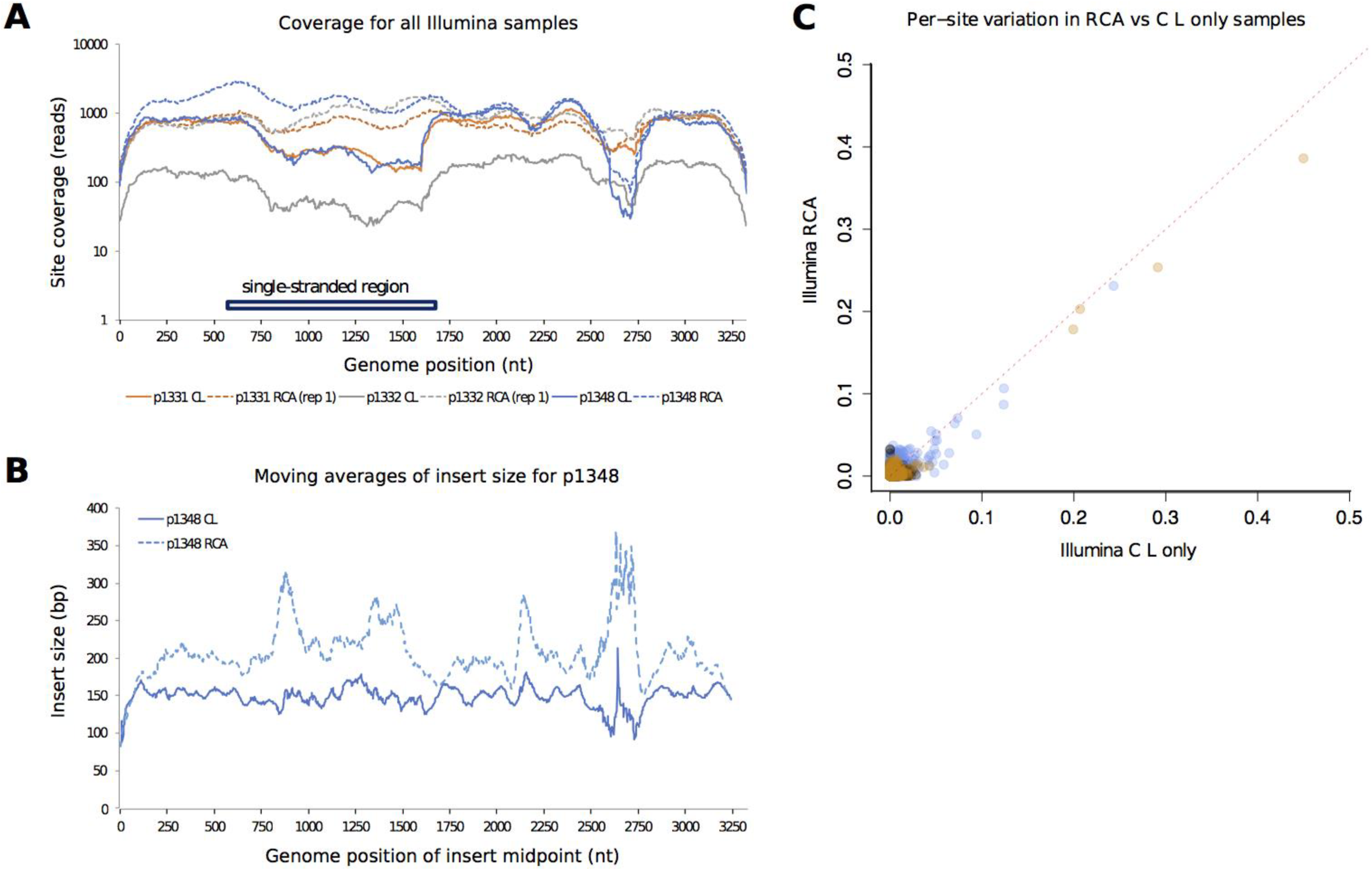
Comparison between HBV sequence coverage and diversity in Illumina sequences generated by completion/ligation (CL) alone, versus CL followed by Phi 29 rolling circle amplification (RCA). A: Read depth across the length of the HBV genome for samples 1331, 1332 and 1348; B: Average insert size across the HBV genome for sample 1348; C: Variation detected in sequences based on CL alone, vs. CL + RCA. Each point represents a genome position with read depth >100. For each of these positions, variation is measured as the proportion of non-consensus base calls, and plotted for both sample types. The red dotted line indicates y=x. In all plots points are coloured by patient as follows: 1331=orange, 1332=grey, 1348=blue.

To investigate the possible effects of RCA on the representation of within-sample diversity, we compared variant frequencies between CL and CL+RCA. Only 2% of sites had variants at frequency > 0.01 and there appeared to be a consistent reduction in estimated frequency in RCA compared with CL alone (Fig 2C).

### Completion ligation (CL) + rolling circle amplification (RCA) facilitates Nanopore sequencing of full-length HBV genomes

We used the material generated by RCA for Nanopore sequencing on the MinION (ONT) (Fig 1). HBV reads accounted for 0.6-1.3% of all sequences derived from individual samples (Table 1), and for 2.6% of reads in a mixed sample (1331/1332). The majority of the remainder of reads mapped to the human genome (Suppl Fig 2). The reads included full-length concatemers of the HBV genome reaching up to 16 complete genomes per read, with a median of 1-2 genomes per read (Fig 3A, B). The number of reads passing quality criteria required for downstream analysis (described in the methods section) are shown in Table 1.

**Fig 3:**
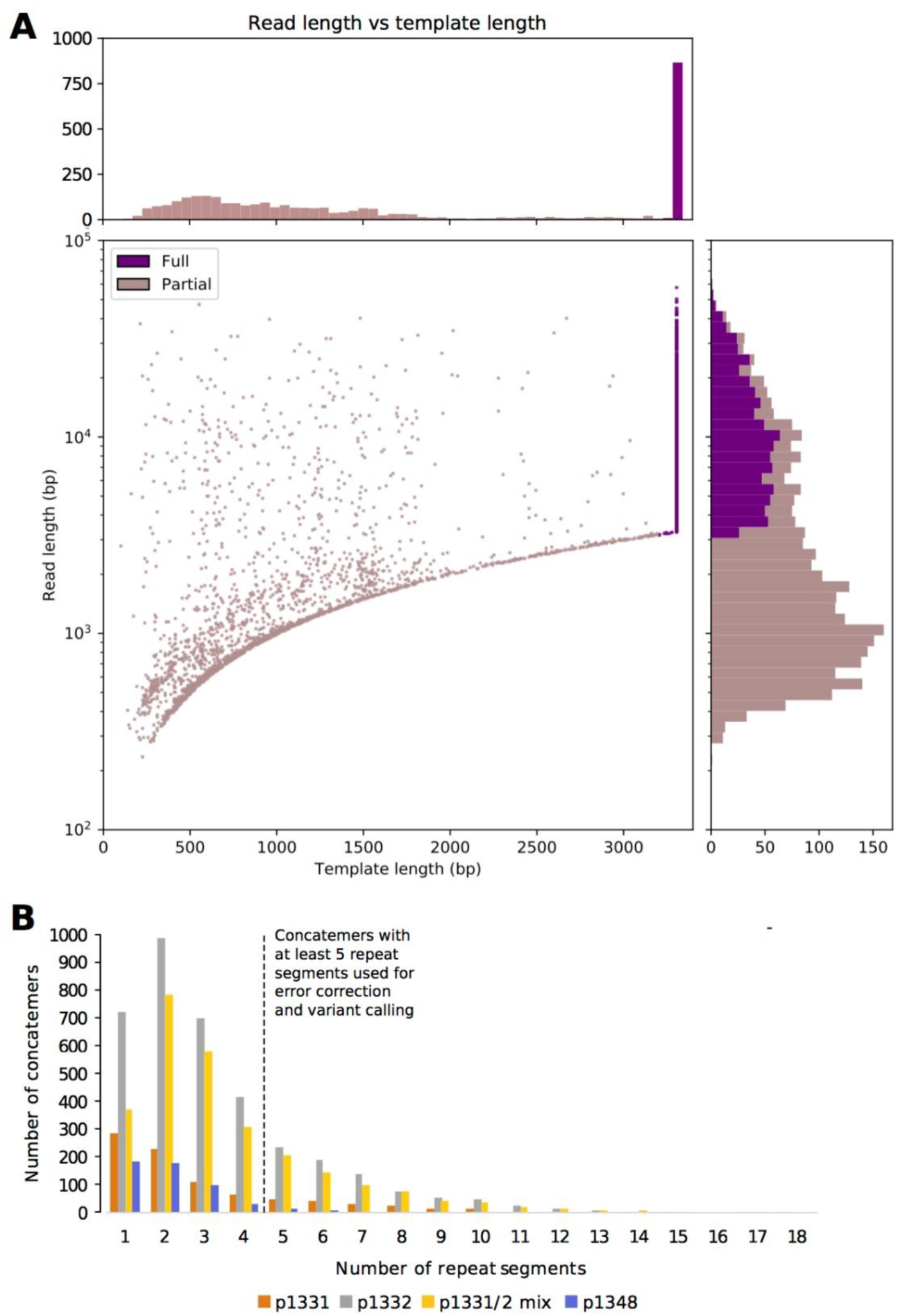
HBV sequence data generated by Nanopore sequencing following completion/ligation (CL) of the genome and rolling circle amplification (RCA). A: Read length and template length of all reads generated from sample 1331. ‘Template length’ refers to the length of the primary alignment of the read, based on a concatenated reference genome. Template length is capped at 3.3kb. Reads with alignments ≥3.2kb in length are considered ‘full length’ concatemers; these are shown in dark purple. B: Plot to show the number of repeat segments in ‘full length’ concatemers. This is equal to the number of segments that a read is chopped into based on the repeated location of an anchor sequence (see methods for details). Reads with ≥ 5 repeat segments will contain ≥ 3 full length copies of the HBV genome. These are taken forward for error correction and further analysis.

### RCA sequencing followed by Nanopore does not produce chimeric sequences in full length genomes

In order to ascertain whether recombination occurred between different viral genomes during RCA or Nanopore sequencing ^11^, we sequenced a mixture of two plasma samples (1331 and 1332, genotypes C and E respectively), producing 3795 reads with a primary mapping to genotype C and 9358 reads with a primary mapping to genotype E. Of these, 148 genotype C and 532 genotype E reads were concatemers containing ≥3 complete genomes and between them they contained 4805 single genome segments. We scored the similarity of each genome segment to the 1331 and 1332 Illumina consensuses at each of 335 sites that differed between the two consensus sequences, classifying genome segments as genotype C or genotype E if they matched the respective consensus at 80% or more of sites (Suppl Fig 3). No concatemers contained segments classified as both C and E. Only 6/4805 genome segments could not be classified in this way, each of which constituted either a short read-end section covering less than eight marker sites, or a low-quality sequence matching variants from both genotypes (Suppl Fig 3). Thus, we found no evidence that the RCA process generates recombined sequences.

### Error correction in Nanopore reads

Among all Nanopore read concatemers with ≥3 complete genome copies, 11.5% of positions differed from the Illumina consensus sequence for that sample. Given Nanopore raw error rates and the observation that the Illumina data contained very few within-host variants, we considered the majority of such differences as likely to be Nanopore sequencing errors. Correcting such errors would allow us to phase true variants into within-sample haplotypes, improving on the information available from Illumina sequencing alone.

As a first step in correcting Nanopore sequencing errors at the whole-read level, we took the consensus of all read segments in each concatemer. Such an approach involves a trade-off between increasing the minimum number of segments per concatemer for inclusion to maximise accuracy, versus increasing the number of reads under consideration to maximise sensitivity for within-sample diversity. To assess error rates we compared corrected sequences with the Illumina consensus, considering only those sites with <1% variation in the Illumina data. For sample 1331, starting with a total of 909 greater-than-genome-length reads, using at least 8 HBV genome copies enabled us to reduce the mean proportion of positions in reads different from the Illumina consensus to 0.51% among 41 reads. Relaxing the criterion to a minimum of 3 genome copies slightly increased the proportion of positions with a consensus call different from Illumina to 0.88%, but allowed us to analyse data from a higher total number of distinct concatemers, n=208 (Suppl Table 1).

In order to reduce the error rate, while using the maximum number of reads, we adopted a refined error correction method. This procedure was based on two assumptions:

i. sequencing errors are randomly distributed across all concatemers, whereas true genetic variants are consistently seen in HBV genomes associated with subset of concatemers (Suppl Fig 4);
ii. unlike true genetic variants, systematic sequencing errors tend to be associated with a particular k-mer in the underlying sequence and so are nearly always strand-specific (note this implies that repeated errors are not only found in homopolymers, as shown in the examples in Suppl Fig 4).

Therefore, to identify variant sites within each dataset, we tested for an association between base and concatemer at each site, analysing plus strand (+) and minus strand (-) concatemers separately and requiring that an association was found in both strands (+ and -) for the site to be considered truly polymorphic. We additionally tested each site for an association between variant and strand, thus sites with significant strand bias were not considered truly polymorphic. We corrected polymorphic sites using the within-concatemer consensus base, whereas sites that failed this test were corrected using the whole-sample consensus base for all concatemers (Fig 4).

**Fig 4:**
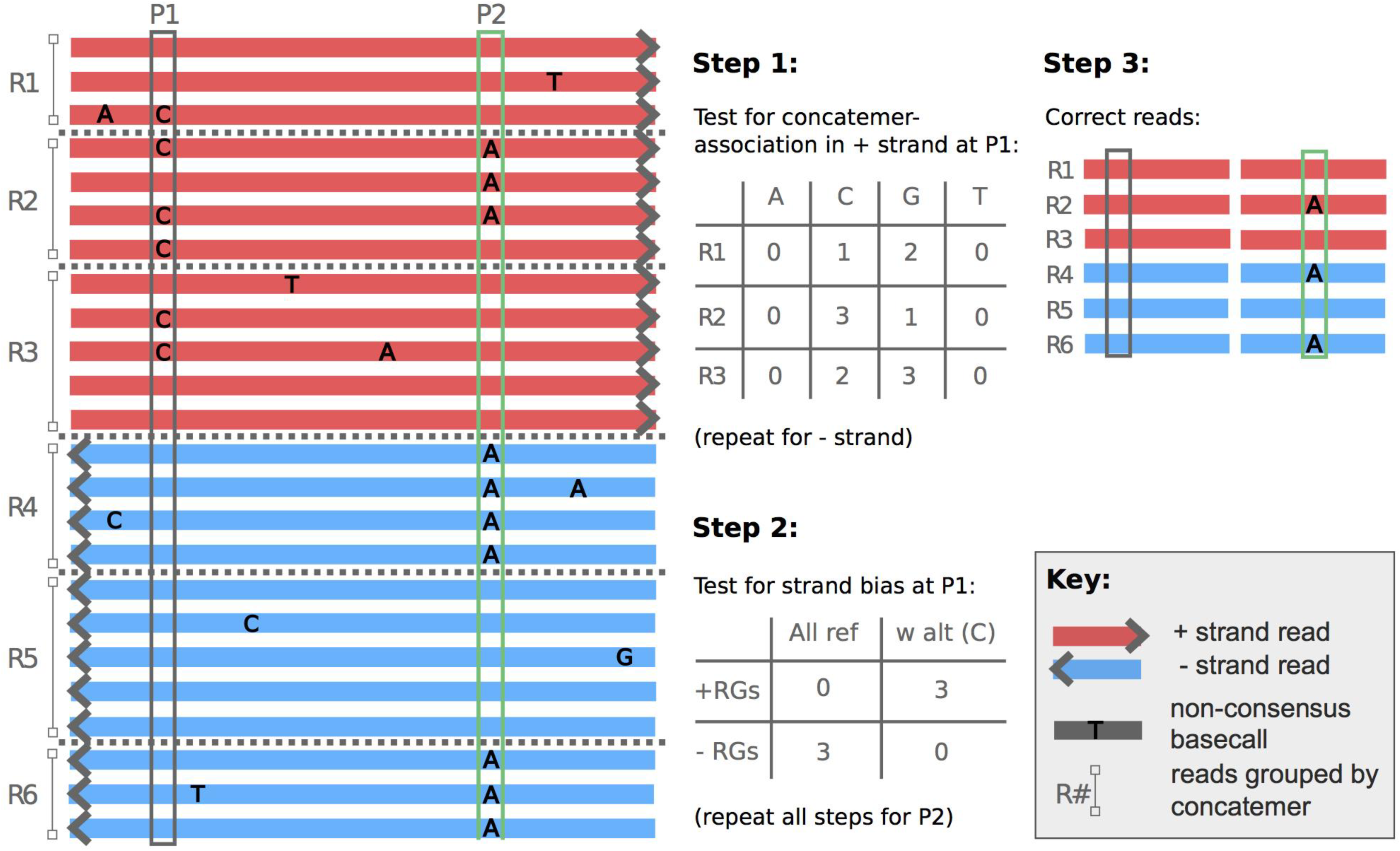
Error correction in Nanopore HBV sequence dataset. A: Schematic to depict the identification and removal of strand-specific basecaller errors. We considered positive (+, red) and negative (-, blue) reads separately. For each set of reads (+ or -), and for each variable site, we tested for an association between base and concatemer using a Fisher’s exact test, such as the contingency table illustrated for + strand concatemers at site P1. A p-value of < 0.01 for both strands was used as the threshold for a significant association. We then tested for strand bias using a Chi-squared test based on the number of + vs - strand read groups (i.e. concatemers) that either contain a basecall for the alternate base or do not. Again, a p-value of < 0.01 was used as the threshold for significance. A lack of concatemer-association, or strand bias, suggest repeated kmer-specific miscalling and that the corrected base should be derived from the consensus across all concatemers. When an association is found for both strand sets and the strand bias is not significant, this implies genome-specific polymorphisms at this site and we set the corrected base to be consensus within each concatemer.

The final corrected Nanopore sequences, each derived from a concatenation of a single HBV genome, differed from the Illumina-derived consensus at an average of <0.4% of sites for the three samples studied (Table 1). We noted that many of these differences were called as gaps (‘-’) or ambiguous sites (‘N’) in the Nanopore data, so the proportion of sites which had been called as an incorrect base was even lower (Fig 5).

**Fig 5:**
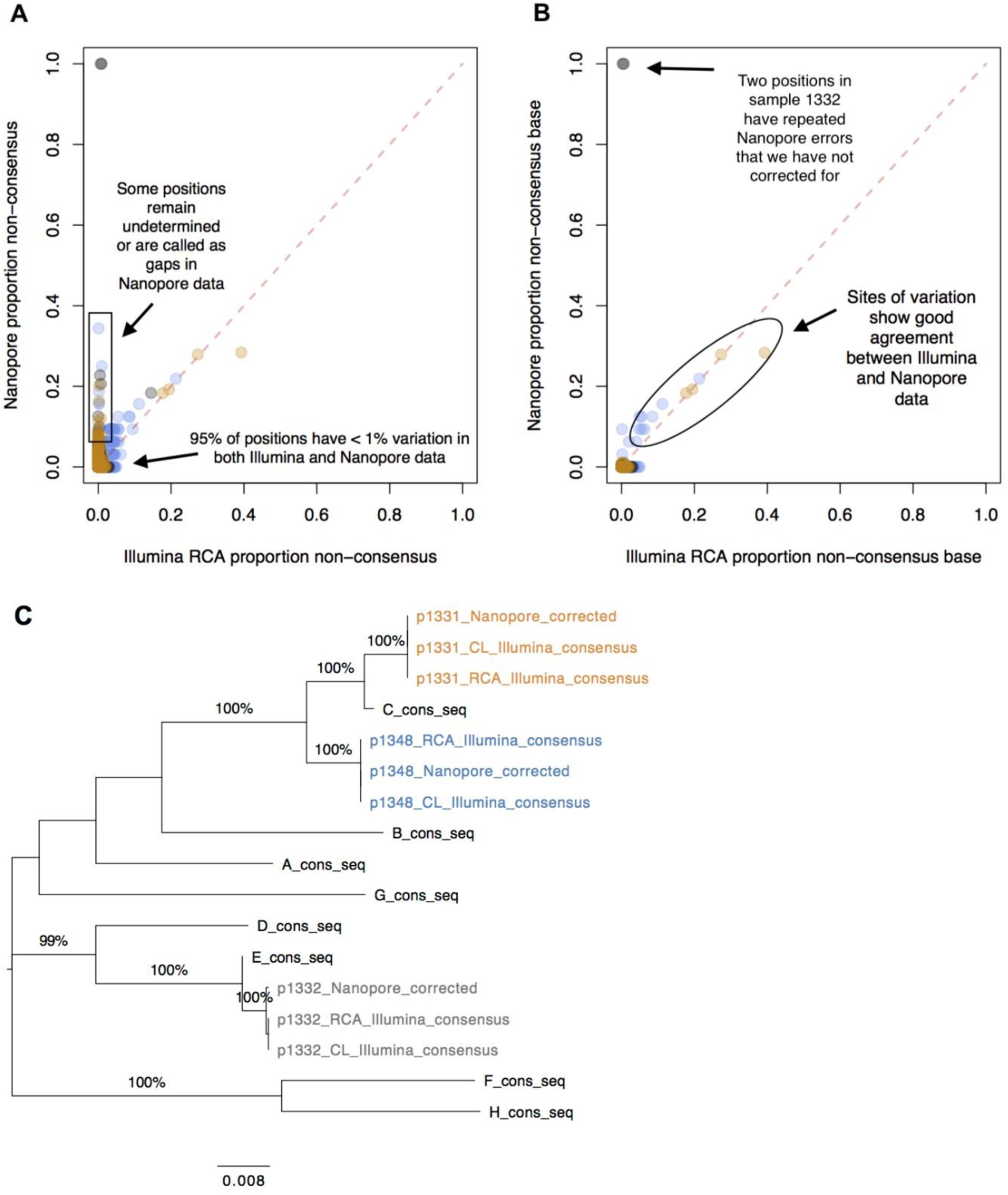
Comparison of HBV sequence data generated by Nanopore vs Illumina platforms, using completion/ligation (CL) and rolling circle amplification (RCA). A: Proportion of non-consensus calls at each position in the genome based on Nanopore (y-axis) vs Illumina (x-axis), for samples 1331 (orange), 1332 (grey) and 1348 (blue). The two sites with 100% variation in Nanopore data are positions 1741-1742 in sample 1332. These lie adjacent to a homopolymer repeat and the high error rate is the result of misalignment when the homopolymer length is miscalled. Positions that are only ever called as ambiguous in the Nanopore data are omitted from this plot (totalling 5 in both 1331 and 1348). Otherwise, sites called as ambiguous (‘N’) or gaps (‘-’) are considered ‘non-consensus’. B: As for panel A, but sites called as ambiguous or gaps are not considered ‘non-consensus’ any more; only alternate bases (A,C,G,T) are included in the ‘non-consensus’ total. C: Phylogenetic tree of consensus sequences for samples 1331 (orange), 1332 (grey) and 1348 (blue) generated by Illumina following CL, Illumina following CL + RCA, and Nanopore following CL + RCA sequencing, together with reference sequences for Genotypes A - H. Bootstrap values ≥80% are indicated. Scale bar shows substitutions per site.

### Detection of true genetic variants in Nanopore data

We then switched our attention to the sites which our Nanopore correction method had highlighted as genuine variants. All variants with >10% frequency in the Illumina RCA data were also detected by the Nanopore method, and frequencies from the two methods showed good concordance (Fig 5). When considering those variants that appeared at >10% frequency in corrected Nanopore concatemers, all were also present at at least 5% prevalence in the Illumina data (Suppl Table 3).

We also used the set of full-length concatemers to derive a within-patient consensus sequence from the Nanopore data. For two out of three samples we found this to be identical to the final consensus sequences for Illumina using CL +/- RCA (excluding 5 sites in each sample which were called as ‘N’s in the Nanopore consensus) (Fig 5C). In the third case, the Nanopore consensus differed at just two sites, located next to a homopolymer (GGGGG).

A primary advantage that Nanopore (long-read data) offers over Illumina (short-read data) is the confident generation of full-length haplotypes, providing insights into the epistatic interactions between polymorphisms at different loci. This is illustrated by quantifying the proportion of genomes derived from Nanopore data that represent a specific haplotype, characterised by combinations of multiple polymorphisms (Fig 6). For example, we were able to identify linkage between two mutations in sample 1348, spaced 510 bp apart in 5/33 whole genome haplotypes (at sites nt 915 and nt 1425, Suppl table 3). Comparing this to Illumina data, the same polymorphisms are detected at similar frequencies but cannot be assigned to a single haplotype in combination. Thus, accurate haplotyping with Nanopore facilitates improved insight into within-host population structure.

**Fig 6:**
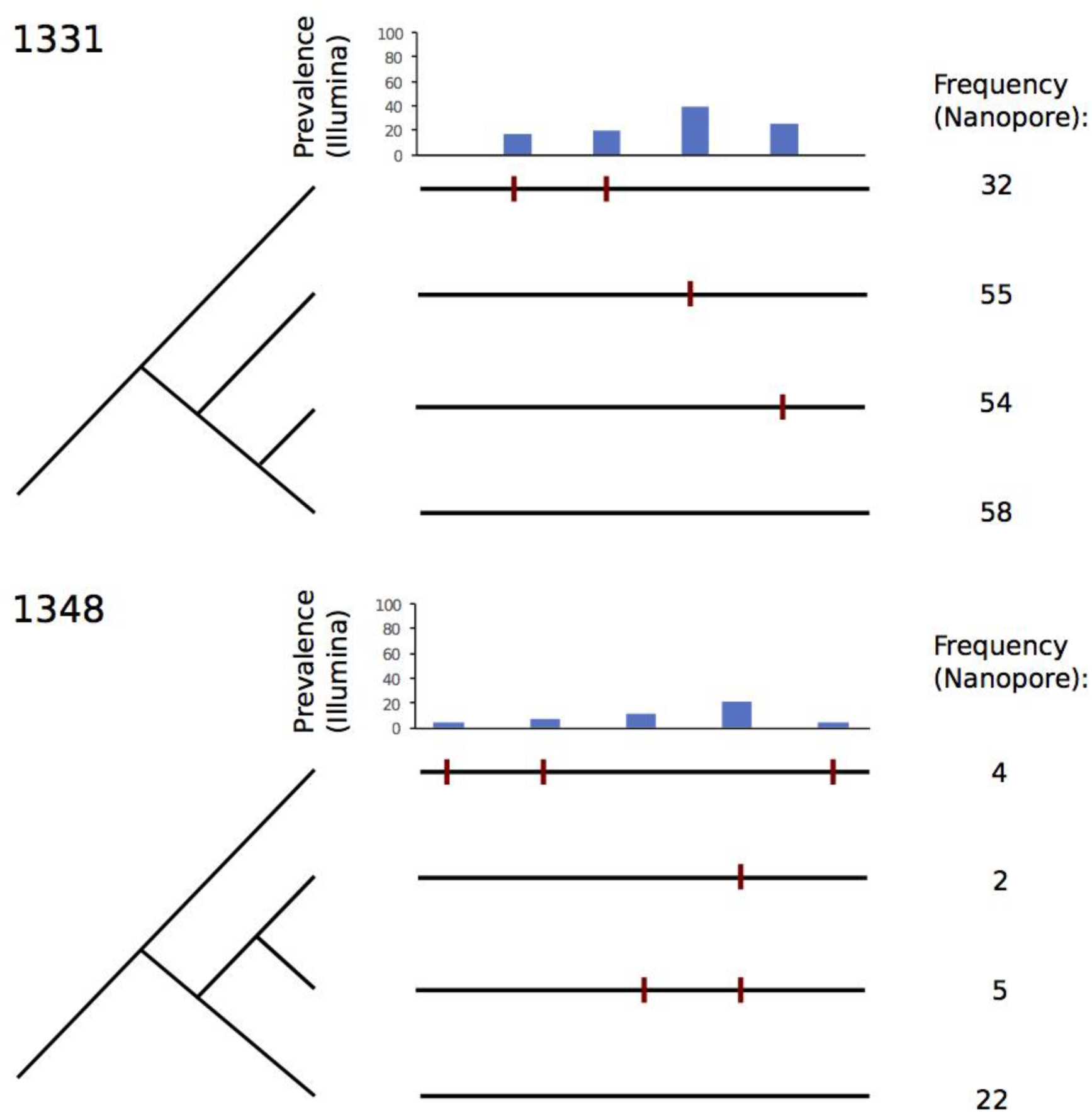
Maximum parsimony trees showing haplotypes called using corrected Nanopore concatemers. For each of samples 1331 and 1348, the high quality variant calls (as listed in Suppl Table 3) were used as a definitive set of variant sites. For each corrected concatemer, the haplotype was called according to the corrected bases at these variant sites. Haplotypes that occurred at >1% frequency within the sample are shown here, with the additional exclusion of one haplotype in sample 1331 that occurred at much lower frequency than those shown (only 3 occurrences) and did not allow for construction of a maximum parsimony tree without homoplasy. Counts of haplotypes are recorded on the left hand side, while the frequency of the variants in the Illumina data is indicated in bar charts along the top of each diagram. Variants (bases differing from the consensus) are indicated with a red bar on the horizontal lines that represent the whole-genome haplotypes.

### Sequence availability

Consensus sequences for our Illumina and Nanopore sequences will be made available on acceptance for publication.

## Discussion

Robust generation of full-length HBV sequence data is an important aspiration for improving approaches to clinical diagnosis (including point-of-care diagnostics and detection of co-infections), patient-stratified management, molecular epidemiology, and long-term development of cure strategies, following precedents set by work in HIV ^17^. However, the unusual biology of the HBV genome, consisting of a small, circular partially dsDNA genome, has represented a significant challenge for whole-genome sequencing. We here demonstrate the use of two different sequencing platforms to generate full length HBV sequences. Illumina approaches allow deep sequencing to determine diversity and detect minor variants, but have the disadvantage of short reads that do not permit the reconstruction of complete viral haplotypes. In contrast, our new Nanopore protocol allows us to gain confidence in the generation of whole HBV genome haplotypes. A comparison of approaches is summarised in Table 2.

**Table 2:**
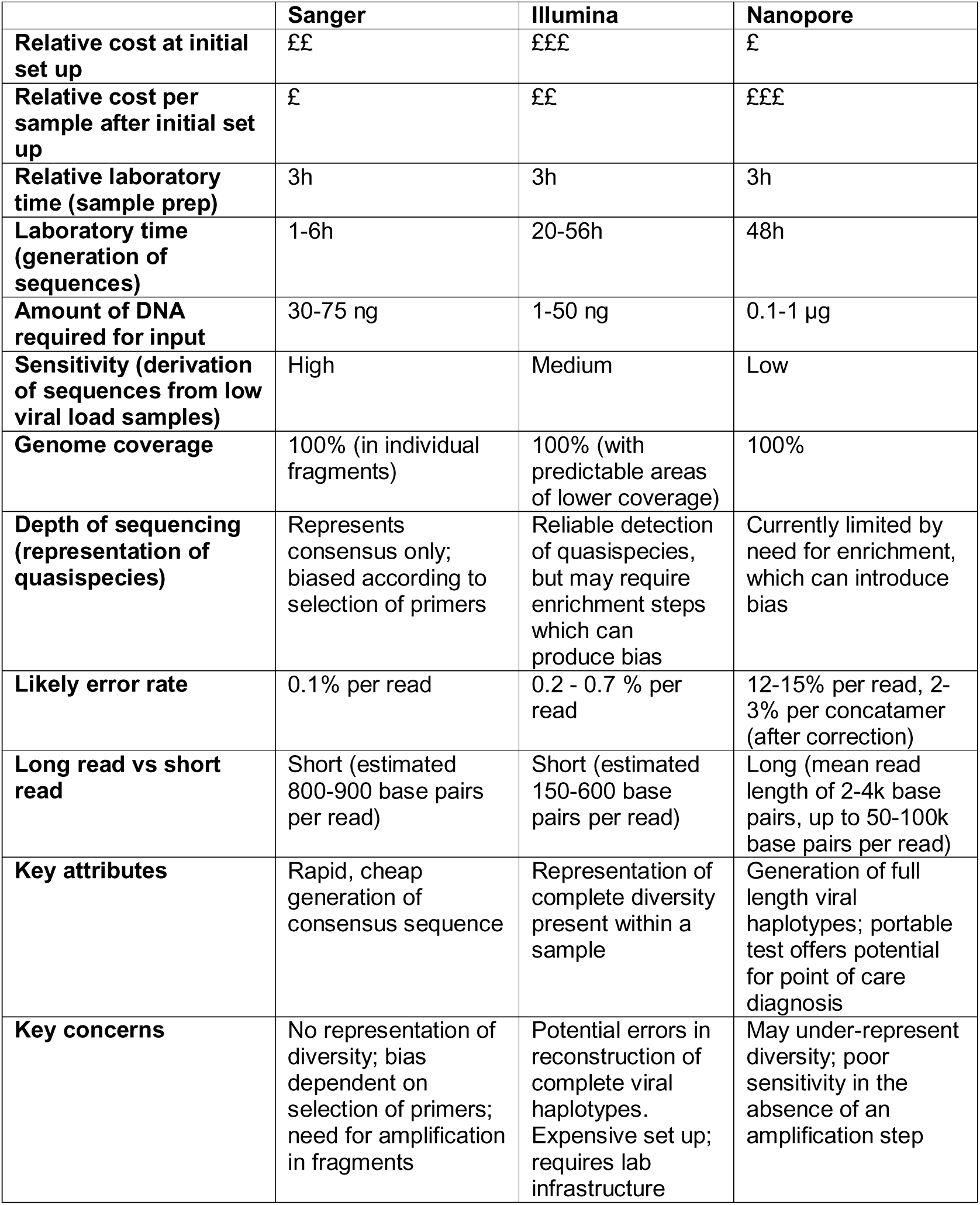
Comparison of three methods of deriving HBV sequence data.

|  | **Sanger** | **Illumina** | **Nanopore** |
| --- | --- | --- | --- |
| **Relative cost at initial set up** | ££ | £££ | £ |
| **Relative cost per sample after initial set up** | £ | ££ | £££ |
| **Relative laboratory time (sample prep)** | 3h | 3h | 3h |
| **Laboratory time (generation of sequences)** | 1-6h | 20-56h | 48h |
| **Amount of DNA required for input** | 30-75 ng | 1-50 ng | 0.1-1 µg |
| **Sensitivity (derivation of sequences from low viral load samples)** | High | Medium | Low |
| **Genome coverage** | 100% (in individual fragments) | 100% (with predictable areas of lower coverage) | 100% |
| **Depth of sequencing (representation of quasispecies)** | Represents consensus only; biased according to selection of primers | Reliable detection of quasispecies, but may require enrichment steps which can produce bias | Currently limited by need for enrichment, which can introduce bias |
| **Likely error rate** | 0.1% per read | 0.2 - 0.7 % per read | 12-15% per read, 2-3% per concatamer (after correction) |
| **Long read vs short read** | Short (estimated 800-900 base pairs per read) | Short (estimated 150-600 base pairs per read) | Long (mean read length of 2-4k base pairs, up to 50-100k base pairs per read) |
| **Key attributes** | Rapid, cheap generation of consensus sequence | Representation of complete diversity present within a sample | Generation of full length viral haplotypes; portable test offers potential for point of care diagnosis |
| **Key concerns** | No representation of diversity; bias dependent on selection of primers; need for amplification in fragments | Potential errors in reconstruction of complete viral haplotypes. Expensive set up; requires lab infrastructure | May under-represent diversity; poor sensitivity in the absence of an amplification step |

The profile of errors in raw Nanopore reads is not often discussed since many users of the technology are primarily interested in obtaining an accurate whole-sample consensus sequence for diagnostic purposes, and error correction tools such as Nanopolish are sufficient for such applications. However the utility of such approaches for the analysis of intra-host diversity is less straightforward and methodological adjustments are required. Our analysis highlights that many errors in raw Nanopore sequence data are kmer-specific, and that these errors are not limited to homopolymer regions. The approach used in this study, using both genome-length concatemers and strand specificity to distinguish kmer-specific errors from genuine diversity, facilitates error correction at the per-read level.

Our results pertain to HBV genotypes C and E, but we believe our methods are likely to be agnostic to genotype, as primers were designed to be complementary to highly conserved regions of the HBV genome 14. Sequencing of a mixed genotype-C/E sample demonstrates that the RCA approach is capable of identifying >1 genotype within a single sample without suggesting recombination, illustrating the reliability of Nanopore long-read data for complete haplotype reconstruction.

The use of RCA to concatenate complete HBV genomes enables error correction at the individual-read level, providing a method with which nanopore-only sequencing can potentially be used to reliably identify and call haplotypes within a single sample. Future optimisation focused on increasing the number of long concatemers will improve the specificity and sensitivity of variant identification and thereby the resolution of low-frequency variants on haplotypes. The methods developed in this study could potentially be applied to study other viruses with small, circular DNA genomes.

In chronic HBV infection, the HBeAg-positive phase of infection is frequently characterised by high viral loads and low viral diversity, as in the samples described here. It has been hypothesised that reduced immune-mediated selection during the HBeAg phase of infection is allowing the unconstrained viral replication of conserved populations ^18,19^, explaining the low diversity we observed in our samples. Marked increases in viral diversity have been described prior to and immediately after HBeAg seroconversion, coinciding with reductions in viral load ^19^. Further work with larger numbers of samples, including those with lower viral loads and HBeAg-negative status, will be of interest in characterising the utility of these different methods for diversity analyses, including identification of specific sequence polymorphisms and determination of within and between host diversity.

Capturing diversity with the Nanopore approach relies on the generation of a good number of concatemers with multiple full-length repetitions. The thresholds chosen to minimise error rate while optimising the amount of sequence data available for analysis may change according to parameters that include viral load, sample diversity, and the question being addressed. Optimisation for lower viral loads is particularly important for the approach to become widely applicable. Broadly speaking, sensitivity can be optimised either through viral enrichment, for example using probe-based selection ^20,21^ and/or by using laboratory approaches that deplete human reads ^22^.

## Methods

### Patients and ethics

We used plasma samples from adults (aged ≥18 years) with chronic HBV infection attending outpatient clinics at Oxford University Hospitals NHS Foundation Trust, a large tertiary referral teaching hospital in the South-East of England. All participants provided signed informed consent for participation. Ethics permission was given by NHS Health Research Authority (Ref. 09/H0604/20). HBV DNA viral loads were obtained from the clinical microbiology laboratory (Cobas ampliprep). We chose samples for sequencing based on their high viral load; all were HBeAg-positive. Blood samples were collected in EDTA. To separate plasma, we centrifuged whole blood at 1800rpm for 10 minutes. We removed the supernatant and stored in aliquots of 0.5-2ml at −80°C. We selected samples of minimum volume 0.5ml and with viral loads of minimum HBV DNA viral load >10^7 IU/ml to optimize successful amplification and sequencing (Table 1).

### Nucleic acid extraction

We extracted total nucleic acid from 500 μl plasma using the NucliSENS magnetic extraction system (bioMérieux) and eluted into 35μl of kit buffer as per the manufacturer’s instructions.

### Completion/ligation (CL) and Phi 29 rolling circle amplification (RCA)

For each sample, we prepared CL reactions in triplicate using previously described methods ^15^. We modified this protocol to maximise the amount of DNA added, by using 6.4 μl extracted DNA plus 3.6 μl reaction mix to obtain a total reaction volume of 10 μl. We retained one reaction for sequencing after undergoing only the CL step, and the other two underwent RCA, using the previously described Phi 29 protocol ^15^. Primer sites are shown in Suppl Fig 5.

### Library preparation and sequencing

For each sample, we used both the product of the CL reaction and of the RCA reaction for library preparation using the Nextera DNA Library Preparation Kit (Illumina) with a modified protocol to account for lower input, based on a previously published method ^23^. We sequenced indexed libraries, consisting of short fragments of PCR-amplified template, on a MiSeq (Illumina) instrument with v3 chemistry for a read length up to 300 nt paired-end.

We used the remaining RCA reaction products, consisting of concatemers of the unfragmented template DNA, for Nanopore sequencing. First, we resolved potential branching generated by RCA by digesting with a T7 endonuclease I (New England Biolabs). We carried out library preparation with a 1D Genomic DNA ligation protocol (Oxford Nanopore Technologies, ONT), and sequenced the samples using R9.4 or R9.5.1 flowcells on a MinION Mk 1B sequencer (ONT).

### Analysis of Illumina data

We demultiplexed paired-end Illumina reads and trimmed low quality bases and adapter sequences (QUASR and Cutadapt software), before removing human reads by mapping to the human reference genome, hg19 using bowtie2. We then used bwa mem to map non-human reads to HBV genotype A-H majority consensus sequences, derived from 4500 whole genomes stored on HBVdb ^24^. We used conventional numbering systems for the HBV genome, starting at the EcoR1 restriction site (G/AATTC, where the first T is nucleotide 1). We re-mapped the same reads using bwa mem to each within-sample majority consensus. In a test of accuracy, consensus genomes were locally aligned to contiguous elements (contigs) assembled ‘de novo’ from the trimmed reads (VICUNA software) and found to match perfectly.

### Analysis of Nanopore sequence data

We basecalled raw Nanopore reads of the RCA-concatemers using ONT’s Albacore versions 2.0.2 (samples 1331 and 1332) and 2.1.10 (sample 1348). We trimmed ‘pass’ reads (those with qscore > 7) using Porechop v.0.2.3 (https://github.com/rrwick/Porechop) to remove adapter sequences. We used Kraken to classify reads ^25^ against on a custom database comprised of the human genome and all complete microbial genomes from RefSeq.

We selected concatemers with ≥3 full-length genomes of ≥2.9 kb, to be taken forward for error correction using a technique similar to that previously described ^11^. In brief, we chopped concatemeric reads (consisting of multiple copies of the same template) into repeating sections and aligned them to form consensus sequences, each representing an error-corrected whole genome haplotype. We identified repeat units in concatemeric reads by chopping reads at an anchor sequence comprising the first 100 bp of the relevant genotype reference; where individual anchor sequences were missed because of poor-quality data we used the distance to the nearest anchor sequence as a guide to form individual genomes. Divided-read sections were remapped with BWA MEM to the HBV genotype reference. Concatemers containing read-sections mapping to both the plus and minus strand were removed (representing a total of 13 concatemers across all three patient samples). To select reads with ≥3 full length genomes we selected concatemers containing ≥5 read-sections, since the first and last section of each concatemer are not guaranteed to be full length.

We applied error correction methods (as described in Fig 4A) to generate a corrected consensus sequence for each group of read-sections corresponding to a single concatemer (and thus a single molecule template). All sites where a non-consensus base appeared at >60% frequency within one or more concatemers were considered sites of potentially genuine genetic variation. We scored and filtered each of these potential variants using the following approach:

1. We conducted a Fisher’s exact test of the association between base and concatemer on + and then - strand concatemers. If either of the resulting p-values were >0.01, we removed the site from the list of variants. We used the two p-values, p1 and p2, to generate a phred-based QUAL score by setting QUAL = −10 * log10(p1*p2), as reported in Suppl Table 3.
2. We calculated a strand bias p-value, by applying a chi squared contingency test to the numbers of + vs - strand concatemers with vs. without observations of the alternate allele. If this p-value was < 0.01 then the potential variant was filtered out.
3. We recorded the allele frequency, AF, calculated as the proportion of base calls across all corrected concatemers that are equal to the most common non-consensus base.

Sites failing either the concatemer-association or strand bias criteria were considered Nanopore errors, and were corrected using the consensus base across all concatemers. Variant sites were corrected using the consensus base within each concatemer. Further filtering based on AF > 0.1 was applied for consistency when comparing Nanopore variant calls with variants at >10% frequency in Illumina. These variants are shown in Suppl Table 3.

Whole-sample consensus Nanopore sequences were derived by taking the most common base at each site, if it was at >40% frequency and was most common in both + and - strand data, or calling the site as an ‘N’ otherwise. The code used for data processing, error correction and variant calling is available on github: https://github.com/hr283/RCAcorrect.

We generated maximum likelihood phylogenetic trees using RaxML with a gamma model of rate heterogeneity and a general time-reversible (GTR) nucleotide substitution model, followed by visualisation in FigTree.

## Acknowledgements

The experimental work described here was funded by the Wellcome Trust (intermediate fellowship to PM grant ref 110110). Core funding to the Wellcome Centre for Human Genetics was provided by the Wellcome Trust (award 203141/Z/16/Z). The work presented here was represented in poster format at the European Association of the Society for the Liver (EASL) International Liver Conference, Paris 2018 ^26^, and at the Nanopore ‘London Calling’ Meeting, London 2018. EB is funded by the Medical Research Council UK, the Oxford NIHR Biomedical Research Centre and is an NIHR Senior Investigator. The views expressed in this article are those of the author and not necessarily those of the NHS, the NIHR, or the Department of Health. We would like to acknowledge the support of the Hepatology clinic at Oxford University Hospitals NHS Foundation Trust for their support in recruitment of patients into research cohorts.

## Author Contributions

ALM, DB, MdC, PCM conceived and designed the project. PCM and EB applied for ethical approval. JBM recruited patients and obtained informed consent; clinical blood samples were processed by AB and CdL. ALM, DB and MdC undertook the RCA, Nanopore and Illumina sequencing work with expert input from PP and RB. JM and ALM generated Sanger sequences. SFL contributed to development of sequencing methods. HER, DB, MAA and ALM analysed the data with oversight from PCM and RB. ALM, HER and PCM wrote the manuscript with input from DB, RB and EB. All authors provided editorial comments, and reviewed and approved the final manuscript.

## Additional Information

We have no conflicts of interest to report.

